# Mitotic chromosome binding predicts transcription factor properties in interphase

**DOI:** 10.1101/404723

**Authors:** Mahé Raccaud, Andrea B. Alber, Elias T. Friman, Harsha Agarwal, Cédric Deluz, Timo Kuhn, J. Christof M. Gebhardt, David M. Suter

**Affiliations:** Institute of Bioengineering, School of Life Sciences, Ecole Polytechnique Fédérale de Lausanne (EPFL), Lausanne, CH-1015, Switzerland; Institute of Biophysics, Ulm University, Albert-Einstein-Allee 11, Ulm 89081, Germany

**Keywords:** transcription factors, mitotic chromosome binding, non-specific DNA binding, specific DNA binding, transcription factor search efficiency, transcription factor occupancy, chromatin accessibility

## Abstract

Mammalian transcription factors (TFs) differ broadly in their nuclear mobility and sequence-specific/non-specific DNA binding affinity. How these properties affect the ability of TFs to occupy their specific binding sites in the genome and modify the epigenetic landscape is unclear. Here we combined live cell quantitative measurements of mitotic chromosome binding (MCB) of 502 TFs, measurements of TF mobility by fluorescence recovery after photobleaching, single molecule imaging of DNA binding in live cells, and genome-wide mapping of TF binding and chromatin accessibility. MCB scaled with interphase properties such as association with DNA-rich compartments, mobility, as well as large differences in genome-wide specific site occupancy that correlated with TF impact on chromatin accessibility. As MCB is largely mediated by electrostatic, non-specific TF-DNA interactions, our data suggests that non-specific DNA binding of TFs enhances their search for specific sites and thereby their impact on the accessible chromatin landscape.

## Introduction

Transcription factors (TFs) are central players in the regulation of gene expression. Each TF binds to specific regulatory sequences to regulate transcription of target genes. The ability of TFs to occupy their specific sites in the genome depends on their nuclear concentration, their ability to search the genome and the chromatin environment of their binding sites. TFs search the genome both by 3D diffusion and by facilitated diffusion using local 1D search mediated by sliding, hopping and intersegment transfer (Dror et al., 2016; Hu et al., 2008; Marklund et al., 2013; Subekti et al., 2017; Takayama and Clore, 2011; Vuzman et al., 2010). These local interactions can strongly modulate search efficiency and mainly depend on transient non-specific protein-DNA association (Berg et al., 1981; Hettich and Gebhardt, 2018; von Hippel and Berg, 1989; von Hippel et al., 1974; Winter et al., 1981), which are essentially mediated by electrostatic interactions (Matthew and Ohlendorf, 1985; Takeda et al., 1986; Kenar et al., 1995; Kalodimos et al., 2004; Barbi and Paillusson, 2013; Vuzman and Levy, 2010; Desjardins et al., 2016; Vo et al., 2017). However, non-specific DNA binding of most TFs remains uncharacterized, and thus to which extent this property impacts genome-wide occupancy of TFs is unknown.

A minority of TFs were shown to physically associate with mitotic chromosomes (Raccaud and Suter, 2017). These interactions can be identified by ChIP-seq on purified mitotic cells and co-localization analysis of TFs with mitotic DNA using fluorescence microscopy. While ChIP-seq essentially identifies sequence-specific DNA binding, fluorescence microscopy allows to quantify the association of TFs with mitotic chromosomes independently of their enrichment on specific genomic sites (Raccaud and Suter, 2017). Both non-specific and specific DNA binding of TFs to mitotic chromosomes have been described. However, the often small number of specifically-bound loci on mitotic chromosomes (Caravaca et al., 2013; Deluz et al., 2016; Festuccia et al., 2018; Kadauke et al., 2012), the mild or null sensitivity to alterations of specific DNA binding properties (Caravaca et al., 2013; Festuccia et al., 2016), and the absence of quantitative relationship between mitotic ChIP-seq datasets and fluorescence microscopy (Festuccia et al., 2018) suggest that the co-localization of TFs with mitotic chromosomes observed by microscopy is mainly due to non-specific DNA interactions. Converging evidence from the literature further corroborates this view. SOX2 and FOXA1 are strongly associated with mitotic chromosomes (Caravaca et al., 2013; Deluz et al., 2016), and these also display high non-specific affinity for DNA in vitro (Sekiya et al., 2009; Soufi et al., 2015). In contrast, OCT4 displays less visible association with mitotic chromosomes (Deluz et al., 2016), and has low non-specific affinity for DNA in vitro (Soufi et al., 2015). Finally, FOXA1 mutants with decreased non-specific DNA affinity but retaining their specificity for the FOXA1 motif also display reduced mitotic chromosome association (Caravaca et al., 2013).

Many TFs binding to mitotic chromosomes also display pioneer properties (Caravaca et al., 2013; Kadauke et al., 2012; Soufi et al., 2012; Zaret, 2014), i.e. they are able to bind and open condensed chromatin regions, allowing them to rewire gene expression programs to mediate cell fate decisions. However, the existence of a common molecular mechanism underlying mitotic chromosome binding and pioneer activity remains uncertain.

Here we measured mitotic chromosome binding (MCB) of 502 mouse TFs in live mouse embryonic stem (ES) cells. We show that MCB is predictive of interphase TF properties such as sub-nuclear localization, mobility, and of the large differences in TF ability to occupy specific genomic sites. We propose that the co-localization of TFs with mitotic chromosomes is a proxy for TF non-specific DNA binding properties, which regulate TF search efficiency for their specific binding sites and thereby their impact on chromatin accessibility.

## Results

### Large-scale quantification of TF association to mitotic chromosomes

To measure mitotic chromosome binding for a large number of TFs, we constructed a doxycycline (dox)-inducible lentiviral vector library of 757 mouse TFs fused to a yellow fluorescent protein (YPet) (Figure 1A). This library was used to generate a corresponding library of mouse embryonic stem (ES) cell lines to quantify TF association to mitotic chromosomes by live cell fluorescence microscopy. One day before imaging, cells were seeded in 96-well plates and treated with dox to induce the expression of TF-YPet fusion proteins. Cells were maintained in ES cell proliferation medium and imaged by wide-field fluorescence microscopy. We used a semi-automated pipeline to detect cells in metaphase, which allows for easy quantification of mitotic chromosome binding since chromosomes are most spatially confined in this phase. Of note, we did not observe any obvious differences in co-localization of TFs with mitotic chromosomes between prophase, metaphase and anaphase. We used the Mitotic Bound Fraction (MBF) as a metric for mitotic chromosome binding, defined as the averaged YPet fluorescence intensity on metaphase chromosomes multiplied by the fraction of cellular volume occupied by DNA (as measured by confocal microscopy, see STAR Methods), divided by the total YPet signal (Figure 1A). The reliability of wide-field fluorescence measurements was confirmed by their correlation with those performed by confocal microscopy (Figure S1A-B, Table S1). In total 502 TFs yielded a sufficiently strong fluorescent signal in metaphase to allow measuring their MBF, and for 94% of these we could measure the MBF in at least 10 cells (see STAR methods). We defined three bins of TFs based on visual inspection of the YPet signal in the area occupied by metaphase chromosomes: depleted (YPet signal lower than in the cytoplasm), intermediate (YPet signal equal to that of the cytoplasm) or enriched (YPet signal higher than in the cytoplasm), which corresponded to MBFs < 16.5%, 16.5-23% and >23%, respectively. 24% of TFs fell in the “depleted” bin, 54% in the “intermediate” bin, and 22% in the “enriched” bin (Figure 1B). Most TFs previously reported to be highly enriched on mitotic chromosomes and present in our library, such as FOXA1 (Caravaca et al., 2013), GATA1 (Kadauke et al., 2012), GATA4 (Caravaca et al., 2013), SOX2 (Deluz et al., 2016; Teves et al., 2016), RUNX2 (Young et al., 2007), ESRRB (Festuccia et al., 2016), RBPJ (Lake et al., 2014) and HNF1b (Verdeguer et al., 2010), fell in the “enriched” category (Table S2), suggesting that C-terminal YPet fusion does generally not perturb mitotic chromosome binding of TFs. We also measured the MBF of a subset of TFs in NIH-3T3 cells, yielding similar results (Figure S1C-D, Table S3), suggesting that the MBF is largely cell type-independent. We then clustered TFs according to their DNA binding domains (DBD) and found that members of some TF families (e.g. homeodomain or tryptophan cluster) were more likely to be enriched on mitotic chromosomes (Figure 1C). In contrast, C2H2 Zinc-finger TFs were mostly excluded from mitotic chromosomes, in line with their mitotic phosphorylation that prevents their association with DNA during M-phase (Dovat et al., 2002; Rizkallah et al., 2011). However, the broad range of MBF within each family suggests that DNA binding domain type does not strictly govern the MBF (Figure 1C), suggesting that other TF characteristics must be involved to regulate mitotic chromosome association.

**Figure 1:**
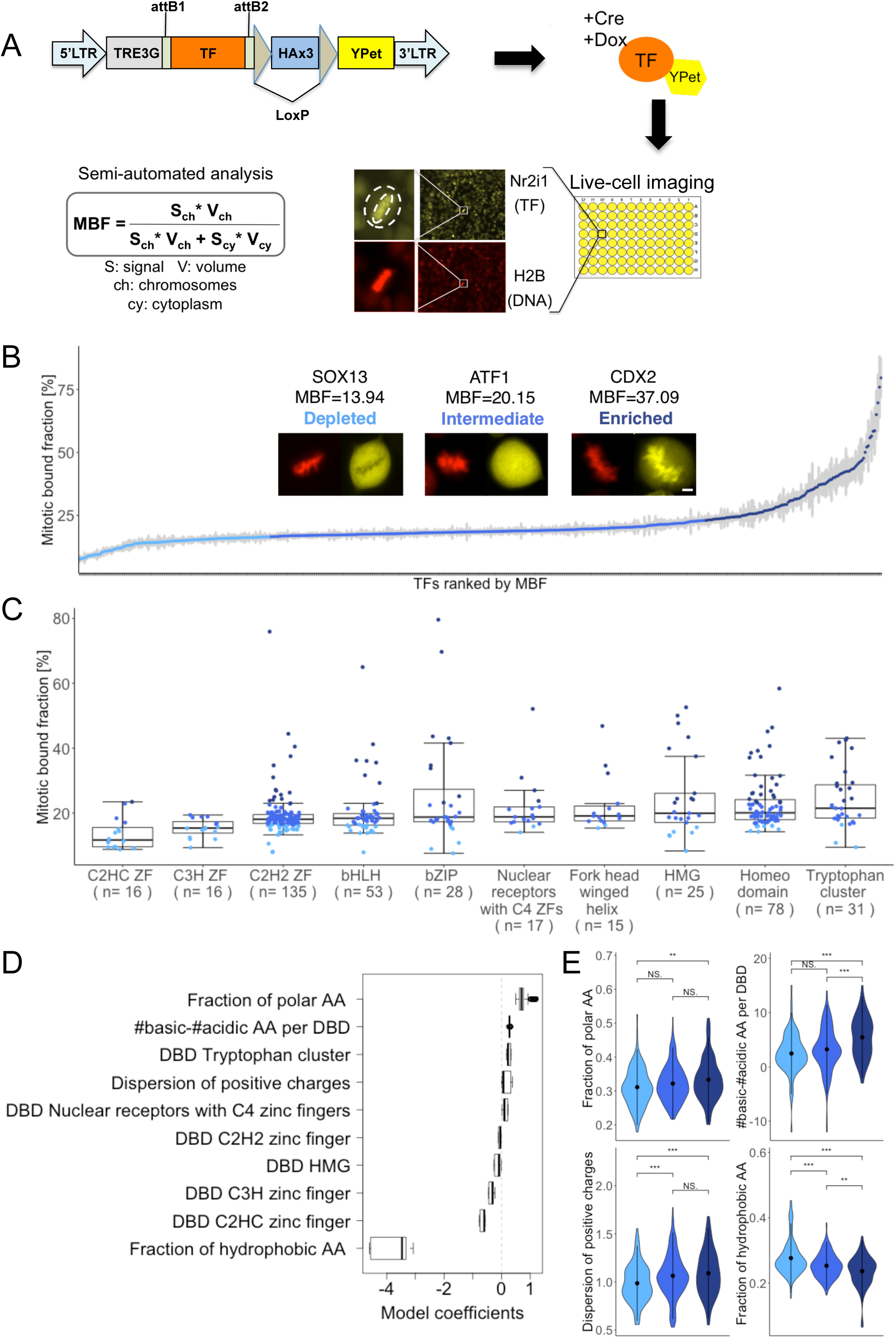
Large scale quantification of mitotic chromosome binding of TFs. **A:** Experimental strategies for generating a lentiviral and ES cell TF-YPet expression library and for quantifying their association to mitotic chromosomes. **B:** Mitotic bound fraction of 502 transcription factors, ranked from lowest to highest and grouped in three bins according to the color code described in the caption. Microscopy inset show representative images of TF localization in metaphase for each category. Scale bar: 3µm. Error bars: SEM. **C:** Mitotic bound fraction of TFs with different types of DNA binding domains. The color code is the same as in panel B. Boxes: intervals between the 25th and 75th percentile and median (horizontal line). Error bars: 1.5-fold the interquartile range or the closest data point when no data point is outside this range. **D:** Parameters recovered by machine learning that impact the MBF and retained in the model for >90% of the runs (n=500). **E:** Violin plots of TF distributions for the fraction of polar amino acids, number of basic residues minus the number of acidic residues, dispersion of positive charges, and fraction of hydrophobic amino acids, grouped in the same categories as in panel B. * p < 0.05; ** p < 0.01; *** p < 0.001; NS. not significant.

### Mitotic chromosome association is correlated with electrostatic properties of TFs independently of active nuclear import

We next used a machine learning algorithm (see STAR methods) using a lasso regularized generalized linear model (Friedman et al., 2010) to uncover TF features based on their amino acid sequences that could explain differences in MBF (Figures 1D and S1E). Briefly, we collected a large number of features extracted from the amino acid sequence of 402 different TFs (see STAR Methods and Table S4) to find those that are correlated with the MBF. We ran the algorithm 500 times on the data to obtain parameters used to predict the MBF of the remaining 100 TFs. The algorithm selects the variables that allow for the prediction of the MBF, while the coefficients of non-predictive parameters are set to zero. Similarly, coefficients of covariates that are correlated among them are set to zero to select a reduced subset of predictive variables. We then kept only parameters with a coefficient >0 in at least 90% of the runs (Figure 1D). As expected, certain types of DBD, such as the tryptophan cluster TF family and the C2HC Zinc fingers were moderately predictive of the MBF. The other parameters most strongly correlated with the MBF were the fraction of polar amino acids, the absolute charge per DBD (number of basic minus number of acidic acidic amino residues), and the dispersion of positive charges (see STAR methods), while the fraction of hydrophobic amino acids was negatively correlated with the MBF (Figure 1D). To determine the impact of these parameters on the whole data set, we compared these physical parameters between TFs falling in the three different bins we defined above (Figure 1B). Although broadly distributed in each bin, all of them were significantly different in TFs enriched on mitotic chromosomes (Figure 1E). The absolute charge per DBD was the most distinctive parameter between TFs enriched on mitotic chromosome (dark blue) versus those that are not (medium and light blue), suggesting that electrostatic interactions play an important role in mitotic chromosome association of TFs.

Since TFs usually harbor one or several nuclear localization signals (NLS), which often consist of a series of positively charged amino acids, we asked whether the impact of positive charges could be confounded by active NLS-mediated transport of TFs to mitotic chromosomes. During mitosis, Ran guanine nucleotide exchange factor (RCC1) associates with mitotic chromosomes and maintains a GTP/GDP gradient between mitotic chromosomes and the cytoplasm (Li and Zheng, 2004), and thus NLS sequences could mediate mitotic chromosome binding through active transport as previously suggested (Lerner et al., 2016; Teves et al., 2016). To discriminate between these two scenarios, we engineered constructs in which we fused YPet to either an NLS sequence or sequences with an equivalent number of positive charges but not predicted to mediate nuclear import. We found that addition of positive charges to the YPet protein were sufficient to increase its co-localization with mitotic chromosomes, independently of nuclear import (Figure S1F). Taken together, these results are in line with previous studies suggesting that i) non-specific DNA binding is essentially mediated by electrostatic protein-DNA interactions, and ii) non-specific DNA binding is largely responsible for the co-localization of TFs with mitotic chromosomes.

### Mitotic chromosome association predicts TF co-localization with DNA in interphase

Since mitotic chromosome association reflects general non-specific DNA binding properties, we next asked whether the MBF scales with TF-DNA co-localization in interphase. To address this question, we used NIH-3T3 cells, which in contrast to ES cells display easily identifiable regions of varying chromatin densities. We stained NIH-3T3 cells with Hoechst and classified the signal in three bins using automatic image thresholding: i) very dense, H3K9me3-enriched heterochromatin regions (Figure 2A); ii) DNA-rich regions; iii) DNA-poor regions (Figure 2B, STAR Methods). We then generated 38 NIH-3T3 cell lines allowing dox-inducible expression of selected TFs fused to YPet spanning a broad range of MBF. After overnight dox treatment and Hoechst staining, we performed two-color confocal microscopy to measure the co-localization of the YPet and Hoechst signals (Figure 2C). We observed a positive correlation between the MBF and the co-localization of TFs with DNA in interphase (Figure 2D and Table S5). We then determined this correlation within the different subnuclear regions we defined, and found the MBF to be positively correlated with enrichment in heterochromatin regions (Figure 2E, Table S5) and to a lesser extent with enrichment in DNA-rich regions (Figure 2F and Table S5). In contrast, we observed a strong, inverse correlation of the MBF with enrichment in DNA-poor regions (Figure 2G and Table S5). Thus, TFs with a high MBF tend to be excluded from DNA-poor regions and are distributed mainly within heterochromatic and DNA-rich regions in interphase, suggesting that non-specific TF-DNA interactions also control interphase TF localization.

**Figure 2:**
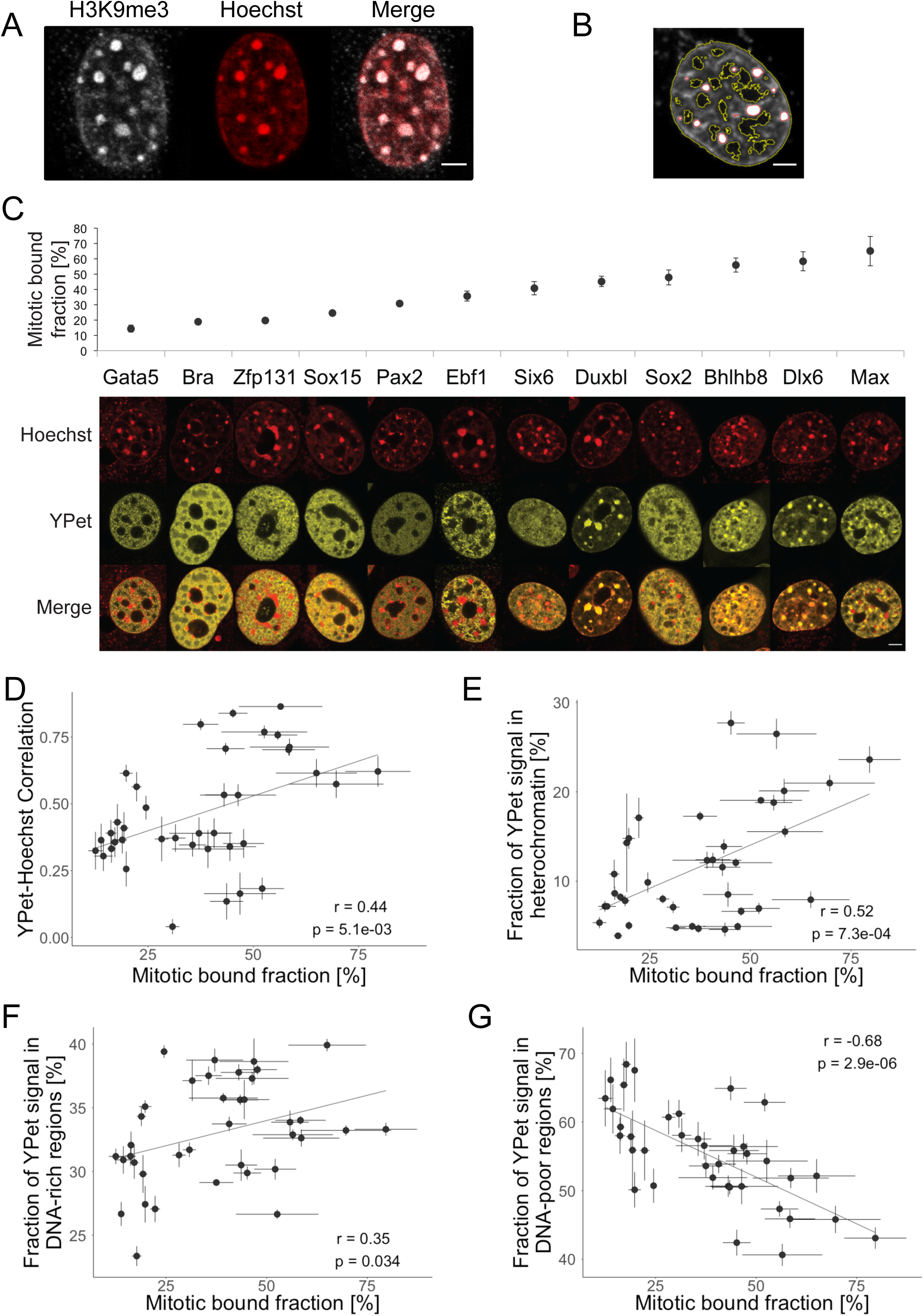
Mitotic chromosome binding is correlated with TF-DNA co-localization in interphase. **A:** Immunofluorescence labeling of H3K9me3 and Hoechst staining of a NIH-3T3 nucleus. Scale bar: 3µm. **B:** Automatic detection of regions displaying different densities of DNA: Heterochromatic (circled in red), DNA-rich (between red and yellow circles), or DNA-poor (circled in yellow). Scale bar: 3µm. **C:** Examples of TF-YPet interphase localization in NIH-3T3 cells, as compared to Hoechst staining and ranked by mitotic bound fraction. Bra: Brachyury. Scale bar: 5µm. **D:** Correlation between the TF-YPet/Hoechst co-localization and the mitotic bound fraction. **E-G:** Correlation between the mitotic bound fraction and the fraction of YPet signal co-localized with heterochromatic regions (E), DNA-rich regions (F), and DNA-poor regions (G). Error bars: SEM.

### Mitotic chromosome association is correlated with TF mobility in mitosis and interphase

We reasoned that co-localization with DNA-rich regions resulting from transient TF-DNA interactions should decrease TF mobility in the cell. To test this hypothesis, we performed fluorescence recovery after photobleaching (FRAP) experiments in ES cells on 15 selected TFs with different MBF in mitotic and interphase cells, as well as FRAP on interphase cells only for 3 TFs excluded from mitotic chromosomes (Figure S2A-B and Table S6). We then used half-time (t_1/2_) of fluorescence recovery as a metric of TF mobility. Mitotic and interphase t_1/2_ of fluorescence recovery displayed a strong positive correlation (Figure 3A and Table S6), and TF mobility was strongly correlated with the MBF (Figure 3B-C). Taken together, this suggests that intrinsic TF properties govern the MBF and TF mobility in both interphase and mitosis.

**Figure 3:**
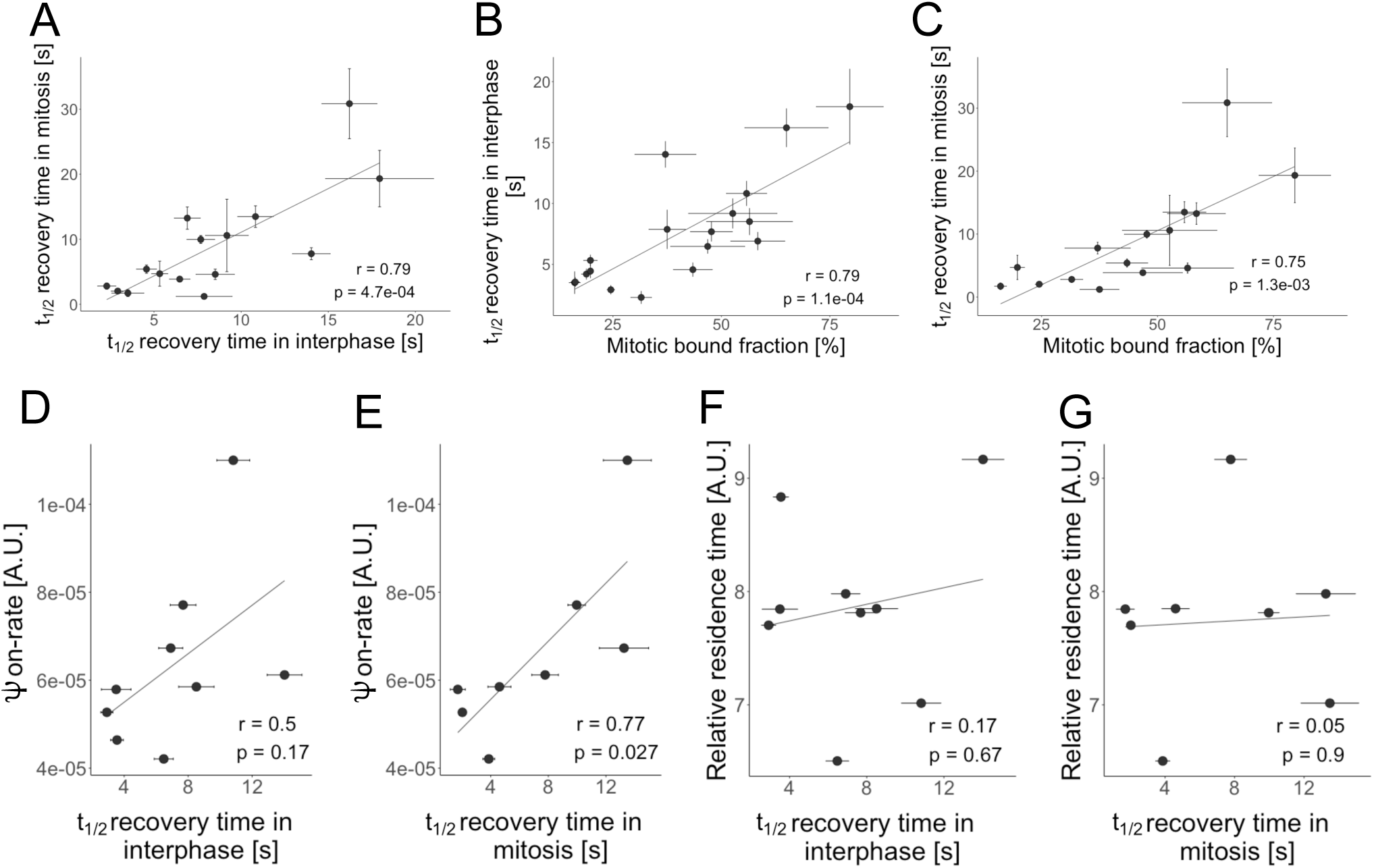
TF mobility correlates with the mitotic bound fraction and interphase ψon-rate. **A:** Correlation between FRAP t_1/2_ recovery of TFs in interphase and mitosis. **B:** Correlation between TF mitotic bound fraction and FRAP t_1/2_ recovery in interphase. **C**: Correlation between TF mitotic bound fraction and FRAP t_1/2_ recovery in mitosis. **D:** Correlation between FRAP t_1/2_ recovery in interphase and the ψon-rate. **E:** Correlation between FRAP t_1/2_ recovery in mitosis and the ψon-rate. **F:** Correlation between FRAP t_1/2_ recovery in interphase and TF relative residence times. **G:** Correlation between FRAP t_1/2_ recovery in mitosis and TF relative residence times. Error bars: SEM.

### TF mobility does not depend on TF size or residence time on specific sites

We next asked which TF properties cause the large differences in TF mobility that we measured. FRAP recovery times depend on 3D diffusion, specific and non-specific DNA binding (Mueller et al., 2013). According to the Stokes-Einstein equation, diffusion scales inversely with the size of molecules. We found a negative correlation between the molecular weight of the different TF-YPet fusions and FRAP t_1/2_ recovery (Figure S2C-D and Table S6), which is the opposite of what would be expected if differences in 3D diffusion were to explain differences in TF mobility. To test whether TFs quantitatively differ in their association with specific DNA sites, we performed single molecule (SM) imaging of TF binding. We generated 9 NIH-3T3 cell lines allowing dox-inducible expression of TFs C-terminally fused to a HaloTag and induced TF expression with low doses of dox shortly before performing SM imaging using highly inclined and laminated optical sheet (HILO) microscopy (Tokunaga et al., 2008) (see STAR Methods). We registered all single molecule binding events according to Hoechst staining intensity (Figure S3A and Table S7), that we used to define three regions similarly as we did for confocal measurements of TF nuclear distribution (see Figure 2A-C). First, we calculated the frequency of DNA binding events lasting >1 second normalized to the total nucleus intensity, which provides a pseudo on-rate that can be compared between different TFs (referred as to ψon-rate hereafter). Second, while our SM acquisition scheme does not allow extracting absolute values for DNA residence times, it enables ranking TFs according to their relative residence times (see STAR Methods). FRAP t_1/2_ recovery times in both interphase and mitosis were correlated with TF ψon-rates (Figure 3D-E, Tables S6 and S7). In contrast, FRAP recovery times were not correlated with their relative residence times (Figure 3F-G, Tables S6 and S7). This suggests that while differences in TF ψon-rates could contribute to differences in TF mobility, differences in 3D diffusion or residence times on specific DNA sites do not.

### Genome-wide occupancy of TFs correlate with ψon-rates but not TF residence times

We next aimed to determine the genome-wide TF occupancy at specific sites for 21 TFs spanning a broad range of MBF, and we also included two FOXA1 mutants that have reduced electrostatic interactions with DNA, decreasing non-specific DNA binding activity without altering their specificity for the FOXA1 DNA motif (Caravaca et al., 2013). As expected, FOXA1 mutants displayed a lower MBF than wild-type FOXA1 (Caravaca et al., 2013) (Figure S3B). For each TF, we generated an NIH-3T3 cell line allowing their dox-inducible expression with three HA tags fused to their C-terminus. We verified that different TFs were expressed at roughly comparable levels upon dox induction (Figure S3C, Table S8), allowing ChIP-seq measurements to provide relative measurements of genome occupancy within a given TF concentration range. We then performed ChIP-seq using the same anti-HA antibody for all TFs and used peak calling to determine the number of specifically bound sites. Remarkably, for the 8 TFs for which we obtained both ChIP-seq and single molecule measurements, TF ψon-rates were strongly correlated with the number of ChIP-seq peaks (Figure 4A, Figure S3D-F, Tables S7 and S8). The offset of the linear fit suggests that a fraction of long-lived SM events are not detected by ChIP-seq. As previously reported, these can result from long-lived, non-specific DNA binding events or indirect TF-DNA interactions (Normanno et al., 2015). In contrast, the number of ChIP-seq peaks was not correlated with TF relative residence times (Figure 4B, Tables S7 and S8). This suggests that the differences in genome-wide occupancy of TFs we observed are mainly due to differences in their association rate but not to longer residence times on specific DNA sites.

**Figure 4:**
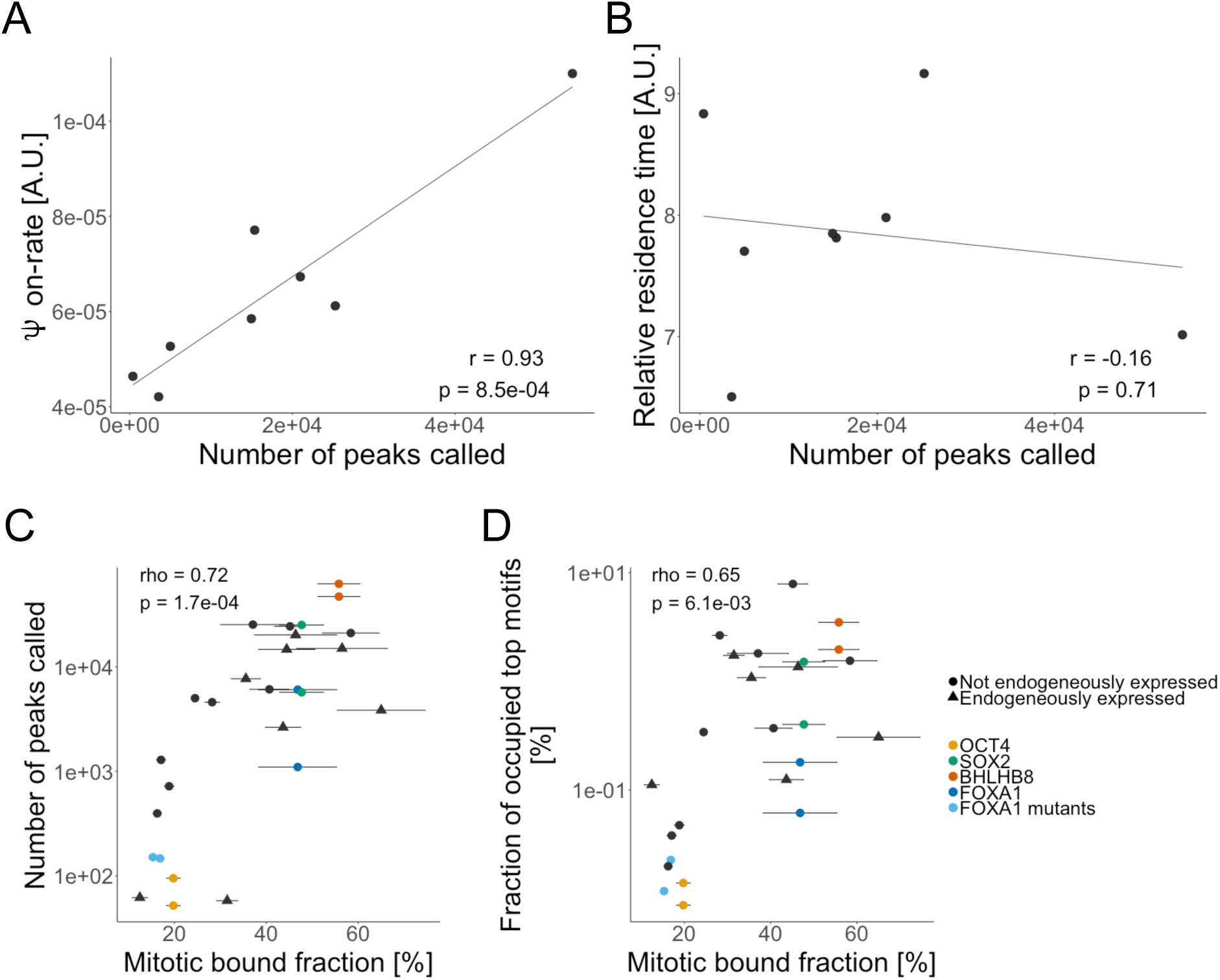
Mitotic chromosome binding predicts genome-wide TF occupancy. **A:** Correlation between the number of ChIP-seq peaks called and the ψon-rate for 8 different TFs. **B:** Correlation between the number of ChIP-seq peaks called and TF relative residence times for 8 different TFs. **C:** Correlation between the mitotic bound fraction and the number of ChIP-seq peaks called. Duplicates are indicated for OCT4 (yellow), SOX2 (green), BHLHB8 (red), and FOXA1 (dark blue). The two FOXA1 mutants are shown in light blue. Triangles: TFs endogenously expressed in NIH-3T3. Circles: TFs not endogenously expressed in NIH-3T3. **D:** Correlation between the mitotic bound fraction and the number of ChIP-seq peaks displaying the most frequently found motif for each TF, normalized over the total number of motif occurrences in the genome. Same color and shape coding as Figure 4C.

### The MBF predicts genome-wide TF occupancy

We used the data obtained from ChIP-seq to compare the relative TF occupancy of specific sites in the genome to their MBF. We first determined the number of ChIP-seq peaks obtained for the 21 TFs, including two biological replicates for OCT4, SOX2, FOXA1 and BHLHB8 to estimate variability in the number of ChIP-seq peaks recovered between two independent experiments, as well as for the two FOXA1 mutants. Remarkably, the MBF and the number of ChIP-seq peaks were strongly correlated, with differences in ChIP-seq peak numbers ranging over three orders of magnitude (Figures 4C and S4A, and Table S8). This correlation was robust to different q-value thresholds (Figure S4B), peak calling algorithms (Figure S4C), and downsampling to equalize the number of reads for each TF (Figure S4D). Very similar results were obtained when using the fraction of ChIP-seq reads in peaks as another metric for TF occupancy (Figure S4E and Table S8), indicating that differences in ChIP-seq peak amplitude are small compared to differences in peak numbers. According to the law of mass action, the rate of formation of a TF-DNA complex scales linearly with its concentration, and therefore the six-fold range in TF expression level (Figure S3C) is unlikely to account for the very large differences in ChIP-seq peak numbers we observed. This relationship was also independent of the presence or absence of endogenous expression of these TFs in NIH-3T3 cells (Figure 4C and Table S8).

We then wondered why some TFs reported to display a high number of ChIP-seq peaks when expressed in their endogenous context displayed orders of magnitude fewer peaks in NIH-3T3 cells. While this may partly be due to differences in TF concentrations, antibodies used, and ChIP-seq protocols, we reasoned that in some cases differences in binding partners could also explain this discrepancy. In particular, OCT4 displayed two orders of magnitude fewer ChIP-seq peaks in NIH-3T3 as compared to ES cells (Liang et al., 2008). As SOX2 and OCT4 form heterodimers, and SOX2 can recruit OCT4 to DNA (Chen et al., 2014; Mistri et al., 2015), we reasoned that OCT4 may have a low intrinsic ability to find its target sites in the absence of a heterodimeric partner. To test this hypothesis, we co-expressed a YPet-SOX2 fusion protein with OCT4-HA in NIH-3T3 and performed ChIP-seq against OCT4. While co-expression of YPet-SOX2 did not alter the expression level of OCT4-HA (Figure S4F), we observed a large increase in the number of peaks (Figure S4G) and fraction of reads in peaks (Figure S4H) of OCT4-HA, suggesting that SOX2 drives OCT4 DNA binding in NIH-3T3 cells.

### The MBF scales with the ability of TFs to find their specific sites in the genome

Importantly, the number of sites bound by each TF may also depend on the number of specific sites available in the genome. To address this question, we quantified the number of occurrences of the most highly enriched (top) motif for the bound sites of each TF (see Table S9 and STAR Methods). We then quantified the fraction of each top motif that is occupied by its respective TF by dividing the number of ChIP-seq peaks containing the top motif (Tables S8 and S9) by the total number of its occurrences in the genome (fraction of occupied motifs or FOM). In addition, we performed ATAC-seq on NIH-3T3 cells to map accessible chromatin regions, allowing us to calculate the FOM in open chromatin regions. The FOM in both the whole (Figure 4D) and accessible genome (Figure S4I) strongly correlated with the MBF (Table S8), suggesting that TFs display large differences in their ability to find their specific sites, independently of the number of sites present in the whole genome or within accessible chromatin regions. Notably, some TFs were not enriched for their known consensus motif from the literature but rather for motifs annotated to other factors. This is not unexpected since TF binding profiles are often context-dependent (Faial et al., 2015; Tsankov et al., 2015), and motif enrichment can also be due to indirect binding or the presence of a motif in proximity of a true binding site. To exclude the possibility that our correlation is biased by indirect binding or incorrect motif identification, we analyzed the FOM at known consensus sites for all factors that were significantly enriched for a motif annotated to the factor itself or a closely related protein (e.g. the SOX3 motif for SOX2 and SOX15, see Table S9 and STAR methods) and found this metric to also positively correlate with the MBF (Figure S4J-K and Table S8). Therefore, our data suggests that the MBF is of strong predictive value for the ability of TFs to occupy their specific sites in the genome at a given TF concentration.

### The broader impact of TFs with a high MBF on chromatin accessibility can be explained by their high genome occupancy

We then asked whether the MBF was predictive of TF ability to modify chromatin accessibility at their binding sites. Since pioneer TFs are suggested to be more capable of targeting closed chromatin regions than non-pioneer TFs (Iwafuchi-Doi, 2018), we first determined if the MBF was correlated with a higher propensity to bind closed chromatin regions. We found no correlation between the MBF and the fraction of TF ChIP-seq peaks in closed chromatin (Figure 5A and Table S8), and a weak positive correlation between the MBF and the FOM in closed chromatin (Figure S5A). We then selected 13 TFs that are not endogenously expressed in NIH-3T3 cells based on a published RNA-seq dataset (Schick et al., 2016) to interrogate their ability to modify chromatin accessibility. For each TF, we performed two biological replicates of ATAC-seq after 48 hours of dox induction in the respective NIH-3T3 cell line. We then determined the number of regions displaying significant changes in their accessibility compared to a control cell line (p<0.05) that coincided with a ChIP-seq peak for that TF (see STAR methods). Overall, the number of regions with changed accessibility coinciding with TF ChIP-seq peaks correlated with the number of TF ChIP-seq peaks (Figure 5B and Table S10). Of note, these regions displayed either significantly increased (Figure S5B) or decreased (Figure S5C) accessibility, and both correlated with the number of TF ChIP-seq peaks. As expected, a large number of genomic sites to which known pioneer TFs such as SOX2 and FOXA1 bound displayed changes in chromatin accessibility (Figure 5B, Figure S5B-C, and Table S10). The impact of TFs on both opening and closing chromatin can be explained by the fact that many of these TFs were reported to function as both activators and repressors (Anderson et al., 2012; Huang et al., 2017; Kawamura et al., 2008; Lemercier et al., 1998; Liang et al., 2008; Liu et al., 2014; Malik et al., 2010). Importantly, these correlations were maintained even when analyzing the same regions for all TFs (defined as all regions with an ATAC-seq peak in at least one sample, see STAR Methods) (Figure S5D-F). To determine the intrinsic ability of each TF to modify chromatin accessibility, we quantified the fraction of sites bound by each TF that displayed significant increase or decrease in chromatin accessibility. Neither of them was correlated with the MBF (Figure 5C-D and Table S10), suggesting that TFs with a high MBF are not more potent on average in altering chromatin accessibility. Therefore, a high MBF is not a signature for an intrinsically higher propensity to alter chromatin accessibility.

**Figure 5:**
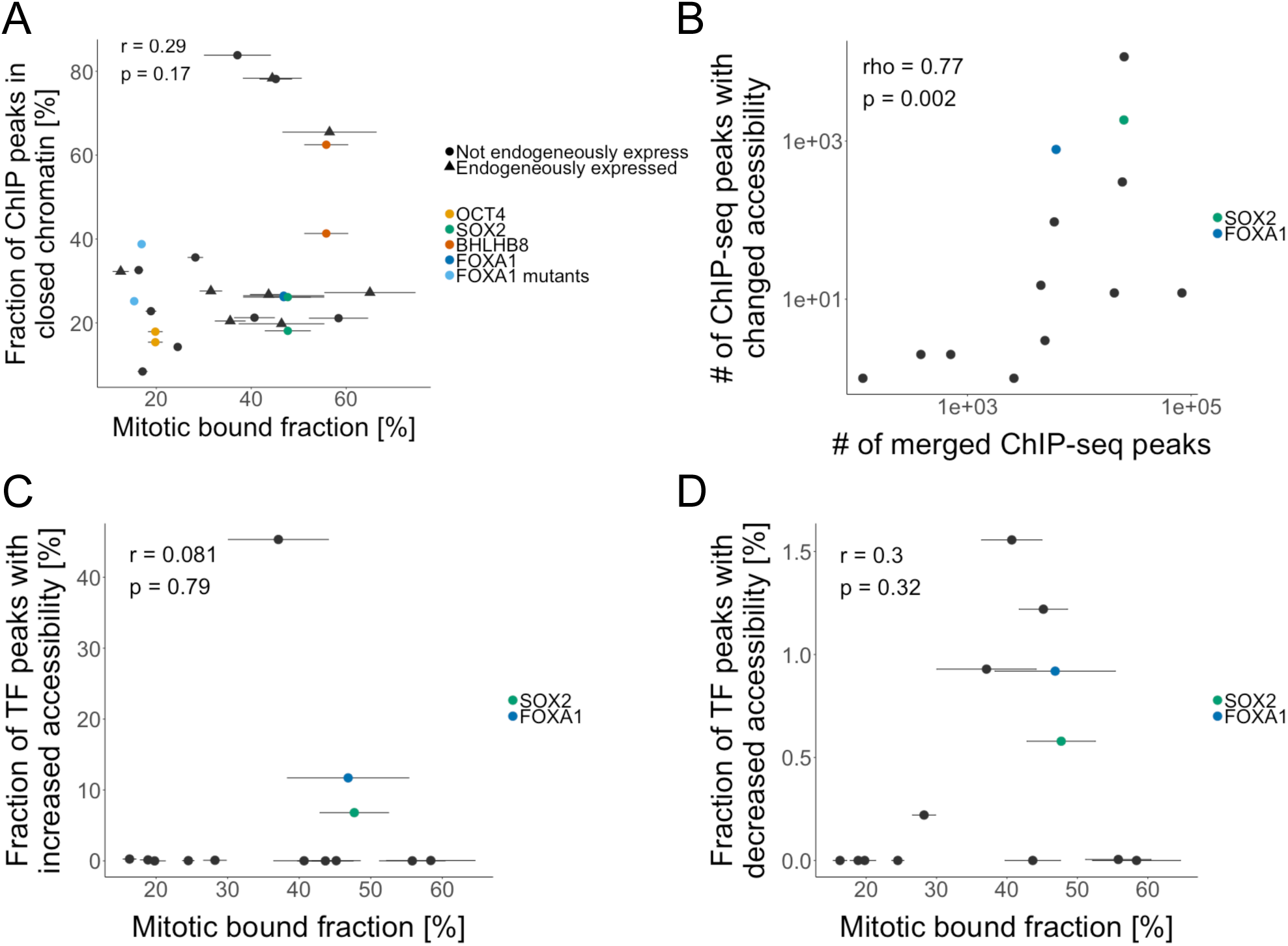
TF capacity to alter chromatin accessibility as a function of genome-wide occupancy and MBF. **A:** Correlation between the mitotic bound fraction and the fraction of peaks in closed chromatin regions (devoid of ATAC-seq signal). Duplicates are indicated for OCT4 (yellow), SOX2 (green), BHLHB8 (red), and FOXA1 (dark blue). The two FOXA1 mutants are shown in light blue. Triangles: TFs endogenously expressed in NIH-3T3. Circles: TFs not endogenously expressed in NIH-3T3. **B:** Correlation between the number of ChIP-seq peaks for each TF and the number of TF ChIP-seq peaks with significant change of chromatin accessibility upon overexpression of the TF. Known pioneers are indicated in green (SOX2) and blue (FOXA1). **C:** Correlation between the mitotic bound fraction and the fraction of regions overlapping a ChIP-seq peak and displaying significant increase in their accessibility upon overexpression of the TF. Same color-coding as Figure 5B. **D:** Correlation between the mitotic bound fraction and the fraction of regions overlapping a ChIP-seq peak and displaying significant decrease in their accessibility upon overexpression of the TF. Same color-coding as Figure 5B. Error bars: SEM.

## Discussion

While it has long been thought that most TFs are excluded from mitotic chromosomes, more recent studies have shown that mitotic chromosome binding of TFs is rather common (Zaidi et al., 2017; Raccaud and Suter, 2017). TFs enriched on mitotic chromosomes were proposed to play a role in maintenance of cell fate by controlling gene reactivation early during mitotic exit (Caravaca et al., 2013; Kadauke et al., 2012; Lake et al., 2014; Zaret, 2014), and the presence of SOX2 and OCT4 at the M-G1 transition was shown to be required for their role in regulating ES cell fate decisions (Deluz et al., 2016; Liu et al., 2017). However, the molecular mechanisms underlying mitotic chromosome association are largely unknown and this property remains uncharacterized for the vast majority of TFs. Here we measured mitotic chromosome association of 502 TFs and found over 100 of them to be enriched on mitotic chromosomes. Our study thus provides a large database of TFs to be mined for their potential role in cell fate maintenance during cell division. While our data indicate that the type of DNA binding domain and electrostatic characteristics of TFs impact mitotic chromosome association of TFs, the *in silico* determination of these properties based on TF amino acid sequence was not sufficient to accurately predict the MBF. Therefore, measuring other parameters that cannot be easily derived from TF amino acid sequence, such as post-translational modifications and three-dimensional structures of TF-DNA contact interfaces is arguably required to allow reliable prediction of the MBF.

We found that quantitative measurements of mitotic chromosome association are of remarkable predictive values for other TF properties. First, differences in the MBF scaled inversely with TF mobility in both interphase and mitosis. This can be explained by differences in non-specific DNA binding properties, which drive mitotic chromosome association and slow down TF mobility in the nucleus. Second, less mobile TFs displayed higher ψon-rates, suggesting that non-specific DNA association increases TF efficiency to search for their target sites through facilitated diffusion. This is also corroborated by the fact that TFs with high and low genome occupancy in our experiments (SOX2 or FOXA1 versus OCT4) display high and low non-specific DNA binding activity in vitro, respectively (Soufi et al., 2015). Third, TFs differed broadly in their ability to occupy specific sites in the genome during interphase, and this property scaled with their association to mitotic chromosomes, even though these two types of measurements were performed in different cell types. Since most of the TFs we studied are not endogenously expressed in NIH-3T3 cells, they are likely to depend essentially on their intrinsic ability to find their specific sites in the genome rather than on the cooperativity with other TFs. Importantly, here we aimed to determine the intrinsic ability of TFs to bind to their sites within a given concentration range. Thus, optimization of TF expression levels in their physiological context may further allow to fine-tune their genome occupancy.

Finally, our data also sheds light on the proposed link between mitotic association and impact on chromatin accessibility. We found that the MBF could predict neither preferential binding to closed chromatin nor the inherent ability to modulate chromatin accessibility. However, the higher number of specific binding sites of TFs with a high MBF was correlated with an increased absolute number of bound sites in closed chromatin regions, as well as a broader impact on chromatin accessibility. Therefore, the on-average higher impact on chromatin accessibility of TFs with a high MBF is mediated by their enhanced ability to occupy specific sites rather than an intrinsically higher ability to alter chromatin accessibility at target sites.

In summary, our findings converge on a model in which non-specific DNA binding properties play a central role in determining TF association to mitotic chromosomes and interphase DNA. We propose that non-specific DNA binding governs TF search efficiency for specific binding sites and thereby their ability to alter the chromatin accessibility landscape (Figure 6). Future studies should allow to further understand the molecular basis of differential non-specific DNA binding activity and how this allows to optimize chromatin scanning to find targets sites in the genome.

**Figure 6:**
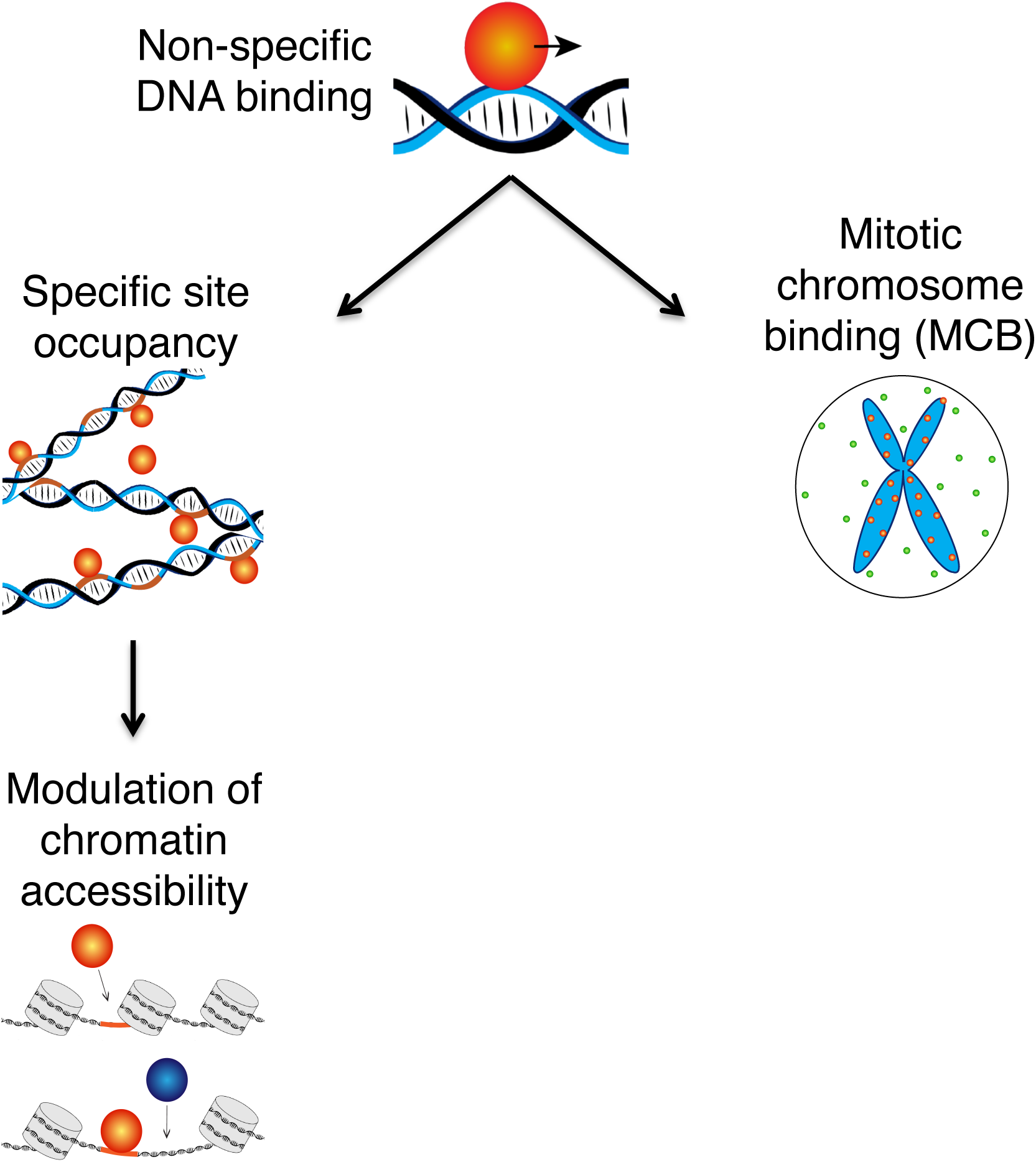
Non-specific DNA interactions drive genome-wide TF occupancy. We propose that non-specific DNA binding regulates mitotic chromosome binding, the search for specific DNA sites and thereby the global impact of TFs on chromatin accessibility.

## Acknowledgments

This work was supported by the Teofilo Rossi di Montelera e di Premuda Foundation advised by CARIGEST SA, and an anonymous donor advised by CARIGEST SA (D.M.S), the German Research Foundation [GE 2631/1-1 to J.C.M.G.], the European Research Council (ERC) under the European Union’s Horizon 2020 Research and Innovation Programme [637987 ChromArch to J.C.M.G.] and the DFG Graduate School of Molecular Medicine at Ulm University (to H.A.). We thank Bart Deplancke for providing the pENTR transcription factor library, Rubina Davtyan and Katharina Soukup for their assistance in analyzing parts of the single molecule data, Vincent Gardeux for his assistance in computing the machine learning algorithm, Bastien Mangeat and Elisa Cora from the EPFL Gene Expression Core Facility (EPFL-GECF) for high throughput sequencing, Fabien Kuttler from the EPFL Biomolecular Screening Facility (EPFL-BSF) and José Artacho from the EPFL Bioimaging and Optics Core Facility (EPFL-BIOP) for assistance in imaging, and Philipp Bucher for critical reading of the manuscript.

## Author contributions

Conceptualization, M.R. and D.M.S.; Methodology, M.R., D.M.S., H.A., and J.C.M.G.; Software, M.R., E.T.F, and T.K.; Formal Analysis, M.R., A.B.A., E.T.F, and T.K; Investigation, M.R., A.B.A., H.A., and C.D.; Resources, D.M.S. and J.C.M.G., Writing – Original Draft, M.R., D.M.S., and J.C.M.G.; Funding Acquisition, D.M.S, and J.C.M.G.; Supervision, D.M.S and J.C.M.G.

## STAR Methods

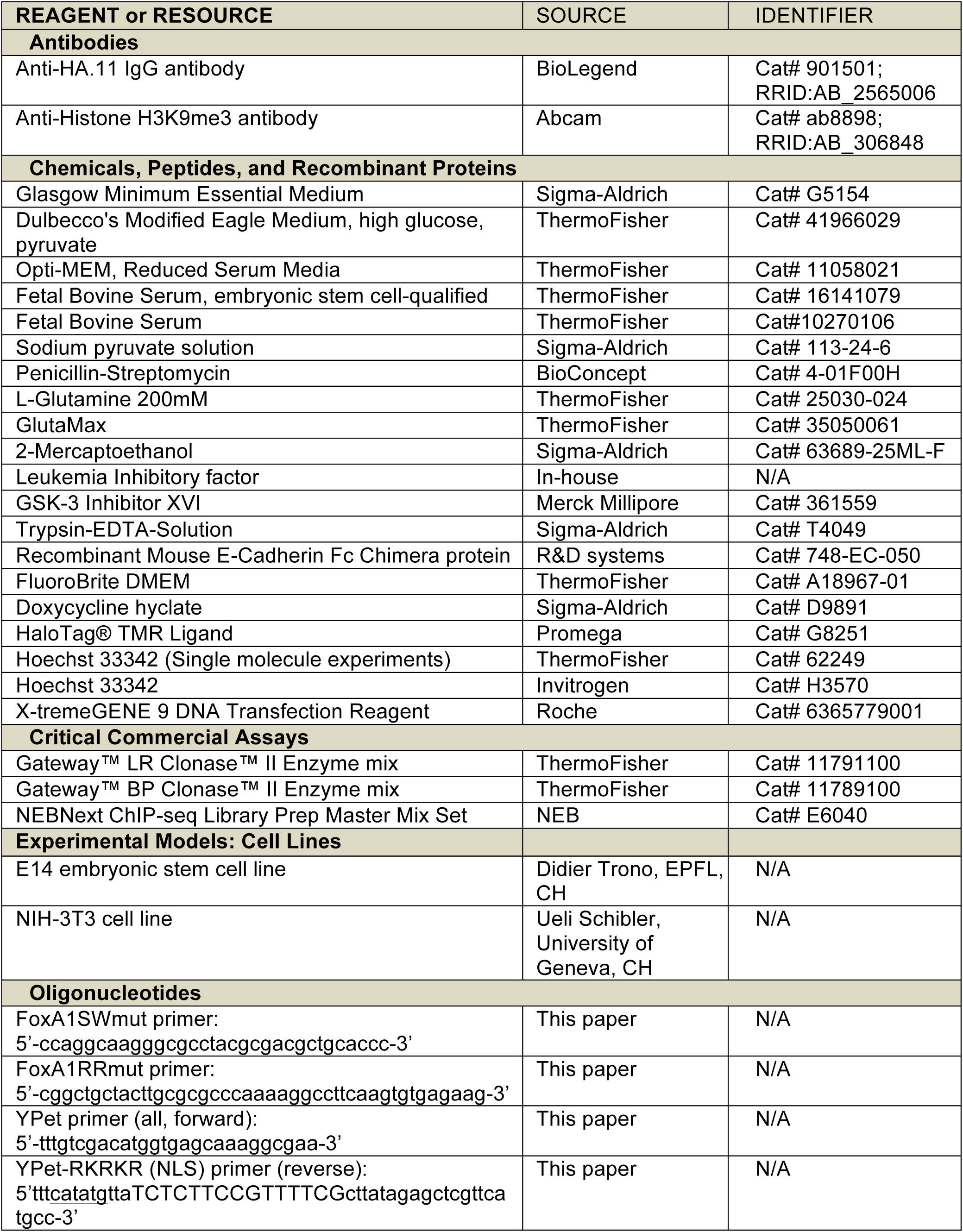
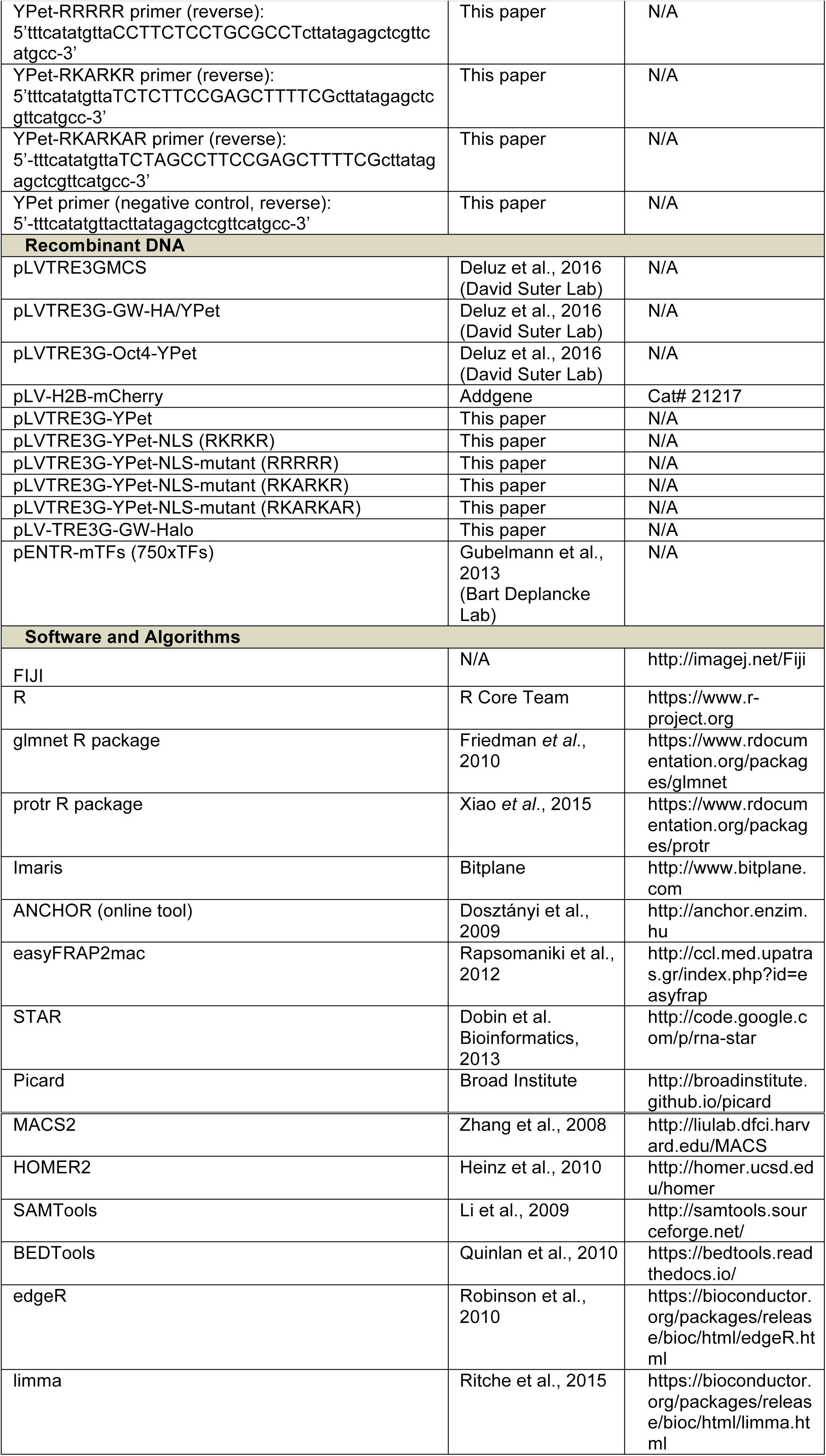
KEY RESOURCES TABLE.

## Contact for Reagent and Resource Sharing

Further information and requests for resources and reagents should be directed to and will be fulfilled by the Lead Contact, David Suter.

## Experimental Model and Subject Details

### Cell lines and culture

The E14 mouse embryonic stem (ES) cell line (provided by Didier Trono, EPFL) was used for all experiments involving ES cells. Cells were routinely cultured at 37°C and 5% CO_2_ in GMEM, supplemented with 10% ES cell-qualified fetal bovine serum, 2 mM sodium pyruvate, 1 % non-essential amino acids, 1% penicillin/streptomycin, 2 mM L-glutamine, 100 µM 2-mercaptoethanol, leukemia inhibitory factor (LIF, concentration not determined, produced by transient transfection of HEK-293T cells and tested for its potential to maintain pluripotency), 3 µM GSK-3 Inhibitor XVI and 0.8 µM PD184352. Cells were grown on 100 mm cell culture dishes coated with 0.1% gelatin, up to confluences of about 70% and split 1:8 to 1:10 every 2-3 days upon trypsinization.

NIH-3T3 cells (provided by Ueli Schibler, University of Geneva) and HEK 293T cells (ATCC) were routinely cultured at 37°C and 5% CO_2_ in DMEM, supplemented with 10% fetal bovine serum and 1% penicillin/streptomycin. Cells were grown in 100 mm cell culture dishes up to a confluence of 90% and split 1:6 every 3-4 days.

For single molecule imaging experiments, we cultured stable cell lines of NIH3T3 cells with different TFs in DMEM supplemented with 10% FBS, 1% Sodium Pyruvate, 1% GlutaMax, 5 µg/ml Blasticidin, 2 µg/ml Puromycin and 1% Penicillin-Streptomycin.

## Method Details

### DNA constructs

A pENTR library containing the coding sequences of 750 mouse transcription factors (TFs) without STOP codon (Gubelmann et al., 2013) was recombined in the doxycycline-inducible expression vector described in (Deluz et al., 2016), (hereafter referred as to pLVTRE3G-GW-HA/YPet). This vector allows for dox-inducible expression of each TF fused either to a YPet (a yellow fluorescent protein) or to three HA tags, depending on the presence or absence of Cre recombinase, respectively. Additionally, the coding sequences of Gbx2, Klf2, Klf4, Klf5, Nanog, Tcf3 and Sox17 were added to the library by PCR amplification from cDNA extracted from either ES cells maintained in the pluripotent state, or from ES cells differentiated for four days by removal of LIF and 2i as described in (Deluz et al., 2016). The PCR amplicons were then inserted in the pDONR221 by BP-Gateway recombination and verified by Sanger sequencing. The pENTR library was then recombined into pLVTRE3G-GW-HA/YPet. The resulting lentiviral vector library was used to generate a corresponding ES library for the large-scale quantification of TF binding to mitotic chromosomes and for fluorescence recovery after photobleaching (FRAP) experiments. In the cases of Sox15-YPet and Hoxd10-YPet, the corresponding E14 cell lines did not yield high enough fluorescence intensity signals to perform FRAP experiments. To circumvent this issue, the coding sequences of Sox15-YPet and Hoxd10-YPet were inserted into the pLVTRE3GMCS backbone (Deluz et al., 2016), which allowed obtaining higher expression levels. This was achieved by removing the Oct4 coding sequence from the pLVTRE3G-Oct4-YPet plasmid (Deluz et al., 2016) using SalI and AscI digestion. The coding sequences of Sox15 and Hoxd10 were then PCR-amplified with primers flanked with SalI and AscI sites, digested with these enzymes, and ligated to SalI/AscI-cut pLVTRE3G-Oct4-YPet.

The two FoxA1 non-specific DNA-binding mutants (Sekiya et al., 2009) were generated by site-directed mutagenesis before recombination in the pLVTRE3G-GW-HA/YPet, by PCR on the FoxA1 pENTR vector with the following primers:

- FoxA1SWmut-F: 5’-ccaggcaagggcgcctacgcgacgctgcaccc-3’
- FoxA1RRmut-F: 5’-cggctgctacttgcgcgcccaaaaggccttcaagtgtgagaag-3’

The constructs for Halo-tagged TFs were generated using restriction cloning, by digesting the pLVTRE3G-GW-HA/YPet with NdeI and AscI to replace the loxP-3xHA-2xSTOP-LoxP-YPet with a NdeI and AscI-cut PCR product of the Halo-Tag coding sequence flanked by NdeI and AscI restriction sites. The selected TFs were then shuttled from the pENTR library into the pLV-TRE3G-GW-Halo by LR-Gateway recombination.

The YPet-NLS and YPet-NLS-mutant constructs were generated by amplification of YPet coding sequence from pLVTRE3G-GW-HA/YPet using 5’-tttgtcgacatggtgagcaaaggcgaa-3’ as forward primer and the following reverse primers to generate the fusion with four different peptides, all bearing five positively charged amino-acids:

- RKRKR (NLS): 5’-ttt<u>catatg</u>ttaTCTCTTCCGTTTTCGcttatagagctcgttcatgcc-3’
- RRRRR: 5’-tttcatatgttaCCTTCTCCTGCGCCTcttatagagctcgttcatgcc-3’
- RKARKR: 5’-tttcatatgttaTCTCTTCCGAGCTTTTCGcttatagagctcgttcatgcc-3’
- RKARKAR: 5’-tttcatatgttaTCTAGCCTTCCGAGCTTTTCGcttatagagctcgttcatgcc-3’
- Negative control, no additional peptide: 5’-tttcatatgttacttatagagctcgttcatgcc-3’

The PCR products were inserted in the pLVTRE3GMCS backbone (Deluz et al, 2016) by restriction cloning using SalI and NdeI restriction sites.

### Lentiviral vector production and generation of stable cell lines

For large-scale quantification of mitotic chromosome binding of TFs, lentiviral vector production was carried out by transfection of HEK 293T cells seeded in 96-well plates with the envelope (PAX2) and packaging (MD2G) constructs together with the lentiviral vector of interest, using the X-tremeGENE 9 DNA Transfection Reagent. Target E14 cells were engineered to constitutively express rtTA3G-IRES-blasticidin, H2B-mCherry and Cre recombinase, referred as to E14 ICC for **I**nducible H2B-m**C**herry **C**re and described in Deluz *et al*., 2016. E14 ICC cells were then used to generate 757 sub-cell lines (among which we were able to grow 753) allowing dox-inducible expression of each transcription factor. To do so, 4000 cells were seeded in 96-well plates and transduced on two consecutive days with 100µl of non-concentrated lentiviral vector particles filtered on MultiScreenHTS 0.45µm filtering plates (Millipore, MSHVS4510). The same transduction protocol was applied to NIH-3T3 ICC cells for quantification of the mitotic bound fraction (MBF) and TF-Hoechst co-localization experiments. Cell lines allowing dox-inducible expression of TF fusions to Halo-Tag for single molecule microscopy, as well as TF fusions to HA tags for ChIP-seq and ATAC-seq experiments were generated by transduction of NIH-3T3 cells constitutively expressing rtTA3G-IRES-blasticidin only.

ES cell lines used for FRAP experiments were generated by transduction of E14 ICC cells with concentrated lentiviral particles generated by calcium phosphate transfection of HEK 293T cells. For all cell lines, selection of transduced cells was performed by addition of the respective antibiotics 48 hours after transduction, and antibiotics were maintained in the cell culture medium throughout passaging. For blasticidin selection, we used 8µg/ml (E14 cells) or 5µg/ml (NIH-3T3 cells); for puromycin selection, we used 2µg/ml (all cell lines).

### Live imaging of TFs and quantification of their mitotic bound fraction

One day before imaging, cells were seeded in black-walled 96-well plates that were either uncoated (NIH-3T3 cells) or coated with E-Cadherin as previously described (Nagaoka et al., 2006), and transgene expression was induced with dox (500 ng/ml). Prior to ES cell imaging, the medium was replaced by E14 imaging medium (FluoroBrite DMEM supplemented with 10% ES cell-qualified fetal bovine serum, 2 mM sodium pyruvate, 1 % non-essential amino acids, 1% penicillin/streptomycin, 2 mM L-glutamine, 100 µM 2-mercaptoethanol, LIF, 3 µM GSK-3 Inhibitor XVI and 0.8 µM PD184352). Prior to NIH-3T3 cell imaging, the medium was replaced by NIH-3T3 imaging medium (FluoroBrite DMEM supplemented with 10 % FBS and 1 % penicillin/streptomycin). Live fluorescence imaging was performed at the Biomolecular Screening Facility of EPFL on an IN Cell Analyzer 2200 apparatus (GE healthcare) with controlled atmosphere (5% CO2) and temperature (37°C) and a 20x magnification objective, using the YFP fluorescence channel for YPet detection, and the TexasRed fluorescence channel for mCherry detection.

For E14 cells, identification of mitotic cells and quantification of the mitotic bound fraction (MBF) were performed using a semi-automated custom pipeline on the CellProfiler software. Briefly, cells in metaphase were automatically discriminated from non-synchronized cells based on shape parameters, and validated by the user. For each confirmed metaphase cell, the selected area was adjusted to precisely define the region containing metaphase chromosomes. This region was then blown-up by 5 pixels and subtracted from a 21 pixel circle drawn around the cell to define the cytoplasmic region. The YPet fluorescence signal was quantified in both regions. For NIH-3T3 cells, regions in the cytoplasm and on the chromosomes were defined manually as described in (Deluz et al., 2016). For both cell lines, the MBF was calculated as:

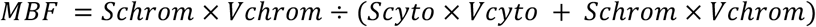

where S is the fluorescence signal and V the average fraction of the volume occupied by either chromosomes or cytoplasm. *Vchrom* (16%) was determined by confocal microscopy on 32 E14 cells. Briefly, an E14 cell line expressing an H2B-mCherry and a cytoplasmic YFP was seeded in Fluorodishes coated with 5 ng/µL recombinant mouse E-cadherin Fc chimera protein at a density of 120,000 cells/cm^2^ and fixed after 24 hours. Mitotic cells were images in 3D stacks on the LSM700 with z steps of 0.496 µm (otherwise same imaging settings as for the NIH-3T3 imaging). The 3D segmentation of the chromosomes and cytoplasm was done using a pipeline developed by the bioimaging core facility of EPFL on the Imaris software. *Vcyto* was determined by *1-Vchrom* (84%). For NIH-3T3, volumes measured for E14 cells were used as a proxy. The MBF was averaged over >10 cells per clone for 94% of the TFs and over 5 to 9 cells for 6% (n=29) of the TFs in E14 cell lines, and over 17 of the 20 NIH-3T3 cell lines, while the 3 others where averaged over 4 to 6 cells.

### DNA-binding domain classification and machine learning analysis

The number of DNA binding domains per TF and their family were extracted from the UniProt database. The DBD families were included in the analysis if present on more than 10 TFs for which we obtained a MBF. TFs with more than one DBD type were included to each of the DBD families. Therefore, in Figure 1C, some TFs are represented in several boxes. Amino acids were classified as in Lee *et al*., 2009, into the following categories: positively charged, aromatic, polar, hydrophobic aliphatic, tiny, bulky, and small amino acids (see Table S4). Additionally, parameters including the sum of the amino acid grand average of hydropaticity (GRAVY) score of the protein, the total number of consecutive positive amino acids, the total number of consecutive neutral amino acids and the total number of consecutive negative amino acids were calculated as previously described (Lee et al., 2009). Most sequence-based parameters were extracted using protr package in R (Xiao et al., 2015). The sum and the fraction of disordered amino acids were evaluated using ANCHOR online tool (Dosztanyi et al., 2009). The dispersion of positive charges was calculated using a sliding window of 5 amino acids to sum the number of arginine and lysine residues and quantified as the variance over the mean of those sums. All absolute counts of amino acids were normalized between 0 and 1. Absence or presence of DBD was annotated with 0 and 1, respectively, for each TF. TFs with missing variables were removed from the analysis. The coefficient for each parameter was calculated using a lasso regularized generalized linear model from “glmnet” package on R (Friedman et al., 2010) on the log of the MBF, and averaged over all the runs. Significant parameters are described as parameters retained by the model in 90% of the runs (n=500).

### Imaging of NIH-3T3 cell lines and co-localization analysis of Hoechst and YPet

NIH-3T3 cells were seeded in FluoroDishes (WPI, FD35-100) at densities of 36,000 cells/cm^2^ 24 h before imaging and treated with 500 ng/ml of doxycycline. Shortly before imaging, cells were incubated with 1.62 µM of Hoechst 33342 for 15 min and washed twice with PBS.

Cells were imaged using a confocal microscope (ZEISS LSM 700 INVERT) with a 63x objective at 37 °C and 5% CO2. Channel settings were as following: EYFP 2.4 % laser power, 700-900 gain, 41.1 µM pinhole; H342 2.6% laser power, 500-700 gain, 39 µM pinhole. Image dimensions: 50.8 × 50.8 µm, 0.05 µm/pixel.

Whole cell signal co-localization was performed by pixel-pixel correlation of the Hoechst and YPet signal images using the R software. Briefly, the Hoechst and YPet images were converted into text images and the correlation score between the two channels was calculated for each cell, using 10 cells per cell line.

The co-localization in different DNA regions was analyzed using an automated image segmentation pipeline in FIJI. Briefly, nuclei were identified and segmented based on the Hoechst signal and 3 regions with high, medium and low Hoechst levels within each nucleus were defined by k-means clustering. Subsequently, the corresponding YPet signal in each of the 3 regions was measured. 10 cells were analyzed per cell line.

### Fluorescence recovery after photobleaching (FRAP) in E14 cell lines

E14 cells were seeded in Fluorodishes coated with 5 ng/µL recombinant mouse or rat E-cadherin Fc chimera protein at densities of 120,000 cells/cm^2^ 24 h before imaging and induced with 500 ng/ml doxycycline.

Cells were imaged using a confocal microscope (ZEISS LSM 700 INVERT) with a 63x objective at 37 °C and 5% CO2. Channel settings: EYFP 2.4% laser power, 700-900 gain, 46.2 µm pinhole. Image dimensions: 200 × 200pixels (19.85 × 19.85 µm).

To image fluorescence recovery after photobleaching, a circular region of interest (ROI) with a diameter of 20 pixels on chromosomes of metaphase cells was selected for bleaching. In addition, two circular control ROIs of the same size were selected, one on the mitotic chromosomes to be used as a non-bleached control and one next to the cell to control for fluctuations of background fluorescence. Cells were first imaged five times with time intervals of 0.38 s to obtain pre-bleach intensity values, and subsequently the selected ROI was bleached for 0.6 s (five iterations) at 100 % laser power. Fluorescence recovery was then imaged for 74 s at intervals of 0.38 s.

The same FRAP experiments were performed in interphase cells (one bleached ROI and one non-bleached control ROI within the nucleus, plus one background ROI next to the cell of interest).

To analyze the recovery time, the mean intensities of the bleached ROI and the two non-bleached control ROIs were measured in all time frames. As mitotic chromosomes tended to move throughout the acquisition, the ROIs on mitotic chromosomes were adjusted manually. The recovery curve of the bleached ROI was normalized based on the intensity values before bleaching and on the two control ROIs. The t_1/2_ recovery time was calculated using easyFRAP2mac (Rapsomaniki et al., 2012) for 10 mitotic and 10 interphase cells.

### ChIP-seq

E14 and NIH-3T3 cells were treated overnight with 500 ng/ml of Doxycycline one day after seeding. ChIP-seq was performed on at least 10^7^ cells as described for asynchronous cells in (Deluz et al., 2016), using anti-HA.11 IgG for all TFs. Libraries were prepared with NEBNext ChIP-seq Library Prep Master Mix Set using insert size selection of 250 bp. Sequencing was performed using 37 nt paired-end reads on an Illumina NextSeq 500. Reads were aligned to the mouse reference genome mm10 using STAR (Dobin et al., 2013) with settings ‘--alignMatesGapMax 2000 --alignIntronMax 1 --alignEndsType EndtoEnd -- outFilterMultimapNmax 1’. Duplicate reads were removed with Picard (Broad Institute) and reads not mapping to chromosomes 1-19, X, or Y were removed. For each sample, peaks were called with MACS2 (Zhang et al., 2008) with settings ‘-f BAMPE -g mm’ (and ‘-q 0.01’ for Figure S4C). Peaks overlapping peaks called for input (non-immunoprecipitated chromatin) from NIH-3T3 cells and ENCODE blacklisted peaks were discarded (ENCODE Project Consortium, 2012). Downsampling of reads (Figure S4E) was done using SAMtools (Li et al., 2009). For HOMER peak calling (Figure S4D), the function findPeaks was used with settings ‘-style factor’ and using Input chromatin as background (for Figure S4D, we added 1 to the number of HOMER-called peaks to avoid zeros). The HOMER2 (Heinz et al., 2010) function annotatePeaks.pl was used with settings ‘-noadj -len 0 -size given’ to count the number of reads in peaks and divided by total aligned reads for each sample to get the fraction of reads in peaks. Motif finding was done using the HOMER2 function findMotifsGenome.pl with settings ‘-size given’. Top motifs were selected as the most significant de novo hit in either the entire peak set for each factor, or in the peaks overlapping (open) or not overlapping (closed) open chromatin based on ATAC-seq peaks in NIH-3T3 (see Table S9). Published motifs were selected by taking the most enriched motif that corresponded to each factor or its TF family (from either de novo or known motif search) (see Table S9). For those factors where no published motif was enriched, JASPAR-annotated motifs were used where possible (DLX1, DLX6, HLF, SIX6, and TEAD1) (Sandelin et al., 2004). In the final analysis, only those motifs with a p-value lower than 0.05 were kept, and factors with top motifs corresponding to “SeqBias” were discarded. The HOMER2 function scanMotifGenomeWide.pl was used to calculate the occurrence of motifs. TFs endogenously expressed in NIH-3T3 were defined as ln(average expression)>2 based on expression data from Schick et al., 2015 (GSE66243). For Figure 4A-B, the average number of ChIP-seq peaks were used for duplicated factors (BHLHB8, FOXA1, SOX2).

### ATAC-seq

NIH-3T3 cells were plated and treated with 500 ng/ml of Doxycycline 48 hours before the experiment. 5*10^4^ cells were collected per replicate and ATAC-seq experiments were performed as described in Buenrostro et al., 2013 using in-house prepared Tn5 transposase (in-house production (Chen et al., 2017)). Two replicates were performed for each TF overexpression sample, and four replicates for control cells expressing only rtTA3G. Sequencing and read alignment was performed as described above for ChIP-seq. To determine regions that were accessible in the NIH-3T3 genome, we performed one ATAC-seq replicate of the parental NIH-3T3 cell line and called peaks as described above for ChIP-seq. ChIP-seq peaks from all TFs analyzed and ATAC-seq peaks from all samples were merged using BEDTools (Quinlan and Hall, 2010) into two separate files and the number of ATAC-seq reads in these peak sets was calculated for each sample using HOMER2. Analysis of differentially abundant regions was done with edgeR (Robinson et al., 2010) and limma (Ritchie et al., 2015), using TMM normalization, comparing each TF overexpression condition to control cells expressing only rtTA3G. For plots displayed in log scale, we added 1 to the number of regions with affected accessibility for each TF. Note that the number of TF peaks represents the number of merged peaks for ChIP-seq duplicated factors.

### Single molecule fluorescence microscopy

#### Sample preparation

Cells were seeded on glass-bottom dishes (Delta T culture dishes, Bioptechs, Pennsylvania, USA) one day before the measurement. To induce the expression of Halo-tagged TFs, 10 ng/ml doxycycline was added 4 hours after seeding the cells. Before imaging, Halo-tagged TFs were labelled with fluorescent SiR ligand (kindly provided by Kai Johnsson, EPFL, Switzerland) according to the HaloTag protocol (Promega). DNA was labelled with 0.3 µg/ml Hoechst 33342. Single molecule imaging was performed in phenol free Opti-MEM at 37 °C for up to 120 min.

#### Image acquisition

Single molecule microscopy was performed on a custom built microscope described previously (Clauß and Popp et al., NAR 2017). Briefly, light of a 405 nm laser (Laser MLD, 200 mW, Cobolt, Solna, Sweden) and a 638 nm laser (IBEAM-SMART-640-S, 150 mW, Toptica, Gräfelfing, Germany) were collimated, combined using a dichroic mirror, controlled by an AOTF (AOTFnC-400.650-TN, AA Optoelectronics, Orsay, France) and used for inclined illumination in a fluorescence microscope (TiE, Nikon, Tokyo, Japan) with a high-NA objective (100x, NA 1.45, Nikon, Tokyo, Japan). Fluorescent light was filtered by a multiband emission filter (F72-866, AHF, Tübingen, Germany) and detected by an EMCCD camera (iXon Ultra DU 897U, Andor, Belfast, UK).

To investigate binding properties of TFs, we used two different illumination schemes: i) continuous movies of cells illuminated with the 638 nm laser to excite SiR were recorded with 50 ms camera integration time, preceded and followed by a snapshot of cells illuminated with the 405 nm laser to excite Hoechst 33342 (50 ms integration time). ii) movies were recorded in which snapshots of cells illuminated for 50 ms with the 638 nm laser were alternated every 550 ms with snapshots of cells illuminated for 50 ms with the 405 nm laser (1.2 s total cycle time).

#### Analysis of bound fractions and chromatin residence times

Detection of individual Halo-TF molecules and identification as bound molecules was performed as previously described (Clauß et al., 2017). In brief, we detected Halo-TF molecules based on their fluorescence intensity above the background level and determined their position using a 2D Gaussian fit. Halo-TF molecules were identified as bound to chromatin when they were detected within a spherical region of 160 nm of diameter for 2 consecutive frames (i.e. for at least 100 ms in illumination scheme i) and for at least 1.2 s in illumination scheme ii)).

We separated chromosomal regions into three classes, bright, intermediate and dark, according to their Hoechst 33342 intensity by means of two user defined intensity thresholds. Subsequent analysis steps were performed separately for each intensity class. We assigned bound molecules to a chromosomal region by comparing their position of first appearance with the preceding Hoechst image. This allowed accounting for cellular movements during long acquisition times in illumination scheme ii).

We determined the fraction of bound Halo-TF molecules per area by dividing the number of molecules identified as bound in illumination scheme i) by the area of the respective Hoechst intensity class and normalized the result by the total number detected of Halo-TF molecules. The fraction of long bound Halo-TF molecules was determined analogously using the respective molecule counts of illumination scheme ii).

We ordered TFs with respect to residence time on chromatin by comparing the average time that Halo-TF molecules spent bound in illumination scheme ii).

### Quantification and Statistical Analysis

Statistical analyses of violin plots of TF distributions (Figure 1E) and on the MBF for NLS mutants (Figure S1F) were performed using Wilcoxon rank-sum test. For plots with linear scales, r- and p-values are based on Pearson correlation (Figures 2D-G, 3A-G, 4A-B, 5A, 5C-D, S1B, S1D, S2C-D, and S3D-F). For plots displayed in log scale, Rho- and p-values are based on Spearman’s rank correlation (Figures 4C-D, 5B, S4B-E, S4I-K, and S5A-F). Note that correlations were calculated using averages for replicates. Duplicate values were not averaged for the FOM of top motifs due to enrichment for different motifs.

